# Mechanism and subunit-dependence of disease-associated *lurcher* mutation in AMPA-type glutamate receptors

**DOI:** 10.64898/2026.11.09.765263

**Authors:** Alexander J. Edwards, Chigdem Arslan, David J. A. Wyllie, Jelena Baranovic

## Abstract

AMPA receptors are glutamate-activated neuroreceptors responsible for the majority of fast excitatory neurotransmission in vertebrate brains. Activity-dependent changes in their functional and trafficking properties underpin cognitive processes such as learning and memory formation. In the past 15 years, a number of mutations within AMPA receptor genes have been identified as disease-causing leading to some form of neurodevelopmental delay. Here, we focused on one such mutation called *lurcher*, resulting from a change of the third Ala in the highly conserved SYTANLAAF motif to Thr. We determined functional consequences of *lurcher* mutation in homomeric AMPA receptors composed of subunits GluA1 or GluA2. In GluA1 subunit, *lurcher* mutation has been identified as a hotspot in a large cohort of patients with neurodevelopmental disorders. We confirm previously established *lurcher* phenotype characterised by slower deactivation and desensitization kinetics as well as more profound block of desensitization compared to the wild-type (WT) homomers. We show that these effects are more pronounced in GluA2 than GluA1 homomers. *Lurcher* phenotypes are attenuated, but still present, in more physiologically abundant heteromeric AMPA receptors composed of two GluA1 *lurcher* and two GluA2 WT subunits (GluA1*^lc^*/GluA2*^WT^*). Schild analysis indicates *lurcher* enhances activity of AMPA receptors by primarily changing the efficiency with which they open their ion channel upon glutamate binding rather than by changing affinity for glutamate. Perampanel, the only widely licensed drug that targets AMPA receptors, is an anti-convulsant that acts as a non-competitive inhibitor. We show it continues to inhibit GluA1*^lc^*/GluA2*^WT^* heteromeric receptors and that its *IC_50_* depends on the auxiliary subunits associated with the heteromers. Specifically, stargazin (TARP γ-2) brings the *IC_50_* for perampanel into the submicromolar range, just like in the WT receptors.

## Introduction

Alpha-amino-3-hydroxy-5-methyl-4-isoxazolepropionic acid (AMPA) receptors are ubiquitous in the vertebrate central nervous system (CNS) where they mediate majority of the excitatory synaptic transmission (Hansen et al., 2021). Activity-dependent changes at synapses in AMPA receptor numbers and subunit composition are critical steps in cognitive processes such as memory formation and learning (Diering C Huganir, 2018; S. J. Liu C Cull-Candy, 2002; S. Ǫ. J. Liu C Cull-Candy, 2000; Nowacka et al., 2026; Penn et al., 2017; Soares et al., 2013). In recent years, an increasing number of mutations in AMPA receptors have been identified as causes of neurodevelopmental disorders (Meagan Collins et al., 2021). Currently, the only widely approved drug that binds to AMPA receptors (and also GluK5-containing kainate receptors (Taniguchi et al., 2022)) is a non-competitive antagonist, perampanel, prescribed to patients suffering from refractory focal-onset and generalized-onset tonic-clonic seizures (Hanada et al., 2011).

AMPA receptors are composed of 4 core subunits, encoded by genes *GRIA1-GRIA4*. In the CNS, the four subunits predominantly combine to form heterodimers (GluA1/GluA2/GluA1/GluA2 or GluA2/GluA3/GluA2/GluA3) or heterotrimers (GluA1/GluA2/GluA2/GluA3) (Herguedas et al., 2019; Lu et al., 2009; Zhao et al., 2019). Homomeric receptors (mainly GluA1 or GluA2(Ǫ) homomers), although less abundant, are also present and perform specific neuronal tasks (Cull-Candy C Farrant, 2021). Subunit GluA2 undergoes developmentally and spatially regulated RNA editing which changes a Gln residue (Ǫ) in the pore into an Arg (R) (Burnashev et al., 1992; Herbrechter et al., 2021; Nutt C Kamboj, 1994; Sommer et al., 1991). The presence of Arg within the pore of an AMPA receptor tetramer greatly reduces Ca^2+^ permeability and overall conductance, while attenuating blocking of the pore by positively charged endogenous polyamines (Bowie C Mayer, 1995; Burnashev et al., 1992; Heiser et al., 2025; Swanson et al., 1997). Although GluA2 is predominantly Ǫ/R-edited in the adult brain, unedited GluA2(Ǫ) subunits more readily assemble into homomeric receptors (Greger et al., 2003). Here, GluA2(Ǫ) was therefore used in homomeric receptors, whereas heteromeric receptors contained GluA2(R). GluA2(Ǫ) homotetramers, in particular, have proven very amenable to biophysical studies and have been the bedrock of structure-function studies of AMPA receptors (Hansen et al., 2021). While the four core subunits are sufficient to form a conducting, glutamate-activated ion channel, native AMPA receptors combine with the so-called auxiliary subunits (Jackson C Nicoll, 2011; Schwenk et al., 2012) (Fig. 1A). The auxiliary subunits primarily associate with AMPA receptors at the level of the transmembrane regions to modulate AMPA receptor trafficking and kinetics (Kamalova C Nakagawa, 2020). Transmembrane AMPA receptor regulatory proteins (TARPs) are the best studied family of AMPA receptor auxiliary subunits, with TARP γ-2 (also known as stargazin) being the most extensively studied member. TARP γ-2 increases AMPA receptor surface expression, slows deactivation, and slows the onset and reduces the extent of desensitization (Priel et al., 2005; Tomita et al., 2005). GluA1/GluA2 AMPA receptor heterodimers associated with γ-2, as examined in this study, are likely present in the mammalian hippocampus and cerebellum (Klugbauer et al., 2000; Lein et al., 2007; Shen et al., 2020; Shen C Limon, 2021), albeit not as the predominant receptor complexes (Scrutton et al., 2026; Yu et al., 2021).

**Figure 1.**
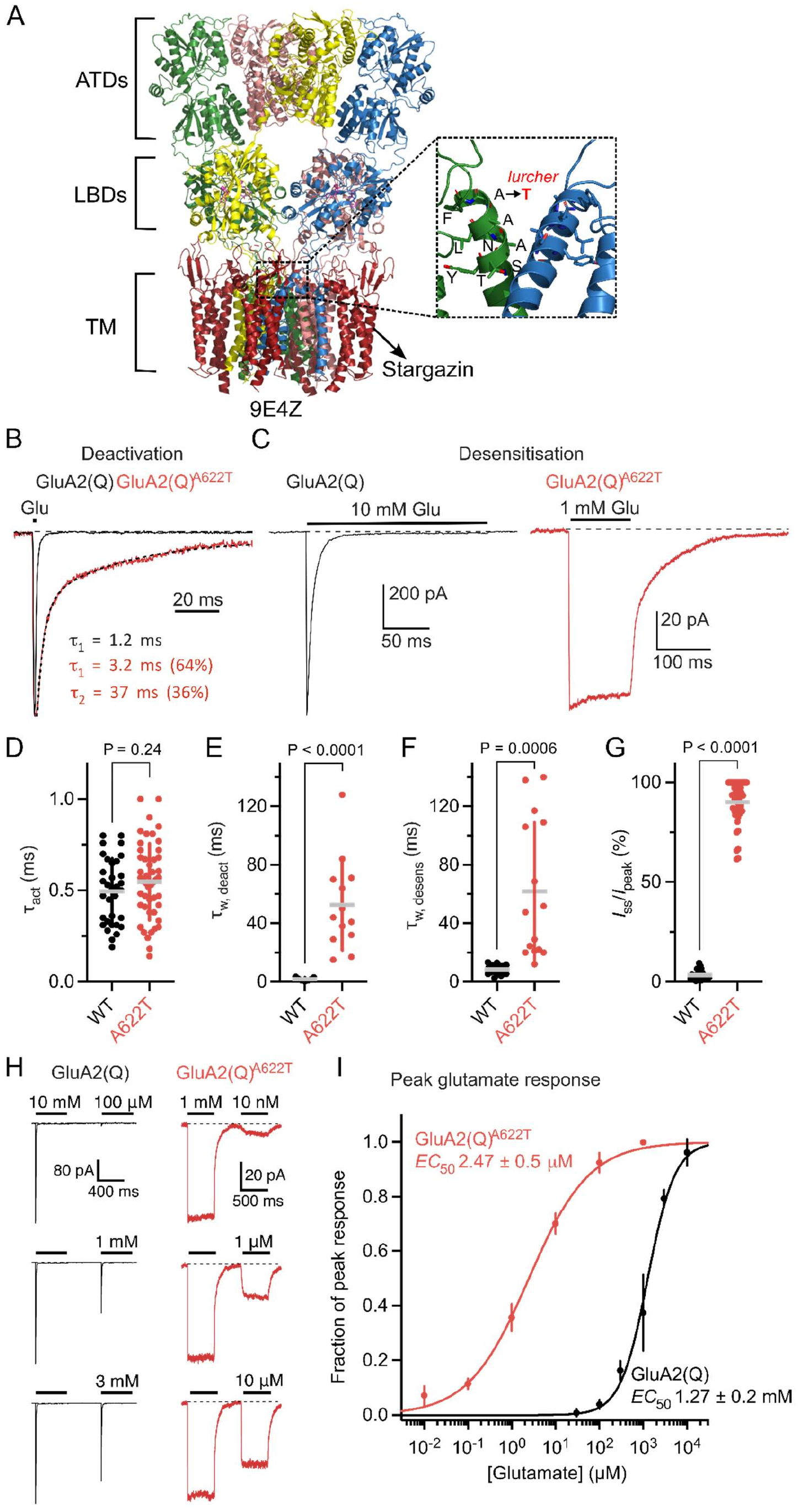
GluA2(Ǫ) wild-type (WT) and GluA2(Ǫ) AC22T lurcher kinetics. (**A**) AMPA receptors are composed of four core subunits (colour coded green, pink, blue and yellow) and, in vivo, associate with auxiliary proteins such as γ-2 (red). Here, all four core subunits are GluA2(Ǫ) (PDB: SE4Z (Carrillo et al., 2025). AMPA receptor domains are indicated by square brackets: amino-terminal domains (ATDs), ligand-binding domains (LBDs) and transmembrane region (TM). The inset shows the region of the highly conserved SYTANLAAF motif, which is also the location of the ion channel gate as well as lurcher (lc) mutation. Pink and yellow subunits and γ-2 have been removed for clarity. **(B)** Representative outside-out patch clamp recordings from GluA2(Ǫ) wild-type (WT) (black trace) and GluA2(Ǫ) AC22T (red trace) receptors in HEK-2S3 cells. To measure deactivation time course, receptors were exposed to saturating glutamate (10 mM for WT and 1 mM for AC22T (lurcher) for 1 ms, at –C0 mV; horizontal black bar). The deactivation decays were fitted with one (WT) or two (lc) exponential components. Traces are scaled to the same amplitude for illustrative purposes. **(C)** Current responses to long, i.e., desensitising applications of glutamate (200 ms for WT and 100 ms for AC22T; 10 mM or 1 mM glutamate respectively, –C0 mV). **(D)** Pooled data points for activation time constants (τ_act_) for GluA2(Ǫ) WT (black spheres) τ_act_ = 0.50 ± 0.18 ms (n = 33) and GluA2(Ǫ) AC22T (red spheres) τ_act_ = 0.55 ± 0.21 (n = 48), P = 0.24, unpaired t-test with Welch’s correction and randomisation test. Grey horizontal lines indicate mean, and vertical bars, the standard deviation (s.d.); circles indicate data points taken from individual patches, with at least three independent transfections per condition. **(E)** As in (D), but for deactivation time constants (τ_w, deact_); GluA2(Ǫ) WT τ_deact_= 1.7 ± 1.1 ms (n = 7) and GluA2(Ǫ) AC22T τ_w, deact_= 52.5 ± 31.0 ms (n = 13), P < 0.0001, unpaired t-test with Welch’s correction and P = 0.0004 with randomisation test. **(F)** As in (D), but for desensitization time constants (τ_w, desens_); GluA2(Ǫ) WT τ_desens_= 8.4 ± 2.7 ms (n = 32) and GluA2(Ǫ) AC22T τ_w, desens_ = C1.8 ± 47.2 ms (n = 15), P = 0.000C, unpaired t-test with Welch’s correction and randomisation test. **(G)** As in (D), but for I_ss_/I_peak_ (%) data points; from GluA2(Ǫ) I_ss_/I_peak_ = 3.0% ± 2.0 (n = 22) and GluA2(Ǫ) AC22T I_ss_/I_peak_ = S0% ± 10 (n = 4S), P < 0.0001, unpaired t-test with Welch’s correction and randomisation test. **(H)** Representative glutamate (black bars) (–C0 mV, 500 ms) current responses for GluA2(Ǫ) WT (black traces) and AC22T (red traces) used for constructing the dose response curves shown in (I). **(I)** Mean dose-response data for WT (black trace) and AC22T (red trace) showing peak glutamate evoked currents (n = 3 patches per concentration). The EC_50_ and n_H_ values were obtained from Hill-type fits and are as follows: GluA2(Ǫ) WT: 1.27 ± 0.2 mM, 1.4 ± 0.4, n = 3; GluA2(Ǫ) AC22T: 2.47 ± 0.5 μM, 0.C ± 0.01, n = 3, P < 0.0001, unpaired t-test with Welch’s correction. Error bars represent s.d; some bars are smaller than the circle size.

Irrespective of their identity, all four AMPA receptor subunits have the same domain architecture: extracellular amino-terminal and ligand-binding domains (ATDs and LBDs, respectively), followed by the transmembrane region (TM) and intracellular C-terminal domain (CTD) (Fig. 1A) (Sobolevsky et al., 2009). Binding of glutamate to LBDs induces conformational changes that result in the opening of the AMPA receptor ion channel gate, located at the extracellular end of the transmembrane helix M3. The ion channel stays open only for a few (5-10) milliseconds and shuts down even with glutamate still bound to the LBDs in a process called desensitization. Channel closure is even faster during deactivation, i.e. if glutamate leaves the LBDs, taking about 1 ms. The channel gate is formed by a highly conserved SYTANLAAF motif present in all ionotropic glutamate receptors (NMDA, kainate, delta and AMPA receptors) (Fig. 1A) (Hansen et al., 2021). The first characterised mutation within this motif was the *lurcher* (*lc*) mutation, based on the mouse delta (δ2) receptor mutant (SYTANLA-‘A to T’-F), which displays a characteristic lurching gait due to swaying of the hind legs (Fig. 1A) (Kohda et al., 2000; Phillips, 1960; Zuo et al., 1997). Since then, mutations in SYTANLAAF motif have been detected in all 4 AMPA receptor subunits in patients with neurodevelopmental disorders (Coombs et al., 2022; Davies et al., 2017; Geisheker et al., 2017; Ismail et al., 2022; Martin et al., 2017; Meagan Collins et al., 2021; Rinaldi et al., 2024; Salpietro et al., 2019; XiangWei et al., 2023).

Here, we set out to understand the mechanism and subunit-dependence behind one such disease-associated mutation in AMPA receptors: change of the third Ala to Thr within the SYTANLAAF motif (C.-M. Chen et al., 2025; Geisheker et al., 2017; Ismail et al., 2022; Rinaldi et al., 2024; XiangWei et al., 2023; Xu et al., 2026). We show that the effects of the *lc* mutation depend on the subunit identity as well as the presence of auxiliary subunits. Our data indicate that *lc* mutation enhances AMPA receptor activity by increasing efficacy of AMPA receptor gating rather than glutamate affinity. The mutation more profoundly blocks and slows desensitization in GluA2 subunits (A622T) than in GluA1 (where the homologous mutation is A636T). Subunit composition also dictates sensitivity to perampanel. In patients carrying the A636T mutation in GluA1, all of whom are heterozygous (de Ligt et al., 2012; Geisheker et al., 2017; Ismail et al., 2022), the mutant subunit will be mostly present in GluA1*^lc^*/GluA2(R)*^WT^* heteromers associated with auxiliary subunits. We show that the presence of the TARP γ-2 auxiliary subunit in the GluA1*^lc^*/GluA2(R)*^WT^* heteromers enhances perampanel potency by reducing *IC_50_* from the micromolar to the nanomolar range.

## Results

### Characterising A622T (*lurcher*) mutation in GluA2(Ǫ) homotetramers

We have characterised *lc* mutation in GluA2 homotetrameric receptors (A622T), observing strongly activating effects: slower rates of deactivation and desensitization, less desensitization and lower *EC_50_* value for glutamate (Fig. 1B-I). While GluA2(Ǫ) WT receptors deactivate with a single-exponential time constant of 1.7 ± 1.1 ms (n = 7), GluA2(Ǫ) A622T mutant shows more complex and slower deactivation kinetics, with τ_deac_ composed of 2 components and an average weighted *τ_w_, _deact_* of 52.5 ± 31.0 ms (n = 13, *P* < 0.0001) (Fig. 1B, E). The *lc* mutation likewise profoundly affects AMPA receptor desensitisation. In contrast to wild-type GluA2(Ǫ) AMPA receptors, which display rapid and nearly complete desensitisation (*τ_desens_* = 8.4 ± 2.7 ms, n = 32; *I_ss_*/*I_peak_* current ratio = 3.0% ± 2.0; n = 22), the weighted time constant for GluA2(Ǫ) A622T is over seven-fold larger (*τ_w, desens_* = 61.8 ± 47.2 ms, n = 15, *P* = 0.0006), and the fractional steady-state current (*I*_ss_) is significantly higher (*I_ss_*/*I_peak_* = 90% ± 10, n = 49, *P* < 0.0001) (Figure 1C, F, G). Since GluA2(Ǫ) A622T did not exhibit sufficient desensitisation, recovery from desensitisation was omitted from this characterisation. We found no change in the time constants for channel activation between WT and A622T receptors (0.50 ± 0.18 ms, n = 33, and 0.55 ± 0.21, n = 48, respectively; *P* = 0.24) (Figure 1D).

Next, we measured the effect of the A622T mutation on the receptor’s sensitivity (*EC_50_*) to glutamate. As the fractional steady-state current (*I*_ss_/*I*_peak_) was in most cases indistinguishable from the maximal peak for the A622T receptors, we used the peak current response evoked at the start of each glutamate application, to generate glutamate dose response curves (Fig. 1H-I). Figure 1H depicts representative current responses from WT (black) and A622T receptors (red) evoked over a range of glutamate concentrations. Each tested glutamate concentration was preceded by a pulse of saturating glutamate which generated a peak response. The test responses were reported as a fraction of the respective peak response. We found that the A622T mutation caused a three orders of magnitude left-shift in the glutamate dose-response when compared to that of GluA2(Ǫ) WT, the glutamate concentrations producing the half-maximal current (*EC*_50_) being 1.27 ± 0.2 mM (n = 3) and 2.47 ± 0.5 μM (n = 3) for WT and A622T (*P* < 0.0001), respectively (Figure 1I).

The observed changes in the kinetics of GluA2(Ǫ) A622T receptors match closely those reported for equivalent mutations in other AMPA receptor homomers: GluA1 (de Ligt et al., 2012; Ismail et al., 2022; Klein C Howe, 2004), GluA3 (Rinaldi et al., 2024), and GluA4 (Zhang et al., 2017). They are all characterised by slower and more complex deactivation and/or desensitization kinetics, reduced extent of desensitization and/or lower *EC*_50_ values for glutamate, with activation time course left unchanged. This suggests all SYTANLA***<u>A</u>***F mutants, with the last Ala mutated into Thr, share the underlying mechanism, irrespective of the subunit identity. We have therefore used GluA2(Ǫ) homotetramers, due to their better expression in HEK cells in our hands, to understand the mechanism behind this mutation.

### *Lurcher* mutation enhances efficacy of AMPA receptor gating rather than glutamate affinity

Fig. 1H-I shows that GluA2(Ǫ) A622T receptors require three orders of magnitude lower concentration of glutamate than GluA2(Ǫ) WT receptors to produce an equivalent half-maximal response (*EC*_50_). The mechanistic basis for this increased sensitivity could reflect a change in glutamate affinity (i.e. binding) and/or change in efficacy with which glutamate opens the ion channel (i.e. its ability to promote conformational change from the resting to the open channel) (Colquhoun, 1998; DEL CASTILLO C KATZ, 1957; STEPHENSON, 1956). To distinguish between the two, we used Schild analysis (SCHILD, 1949; Wyllie C Chen, 2007), which uses the shift in *EC_50_* for an agonist produced by different concentrations of a competitive antagonist to determine the microscopic binding constant for the antagonist. Any large changes in the binding affinity of the competitive antagonist for a mutant would be consistent with the notion that the mutation affects the binding site and correspondingly, agonist affinity. We used NBǪX as a competitive antagonist of GluA2 homotetramers (Sheardown et al., 1990) with a resolved binding site (Coombs et al., 2019; Vega-Gutiérrez et al., 2025) (Fig. 2A). It is a condition of Schild analysis that agonist and antagonist demonstrate mutually exclusive binding to the same site, i.e. engage a majority of the same residues in the binding site. Fig. 2A shows this is indeed the case for glutamate and NBǪX. High resolution crystal structures of GluA2 ligand-binding domains in complex with glutamate (PDB: 2I3W (Armstrong et al., 2006)) and NBǪX (PDB: 6FǪH (Coombs et al., 2019)) show that both ligands engage (i.e. are within 4 Å of) the following residues: Tyr450, Pro478, Leu479, Thr480, Arg485, Leu650, Glu705 and Tyr732 (Fig. 2A). We constructed glutamate dose-response curves for GluA2(Ǫ) WT (Fig. 2B-C) and GluA2(Ǫ) A622T mutant receptors (Fig. 2D-E) in the presence of 0, 3, 10 and 30 μM NBǪX. To block desensitization, cyclothiazide (CTZ; 100 μM) was present throughout, and fractional steady-state (I_ss_) responses were measured to ensure all measurements were made at equilibrium (a condition of Schild analysis). The glutamate dose response curves in both cases showed a near-parallel right shift as NBǪX concentration increased and similar saturating responses, both indicators of competitive antagonism. At 30 μM NBǪX and high glutamate concentrations, the upper asymptote of the concentration-response curve for GluA2(Ǫ) WT receptors could not be fully resolved, as the glutamate concentrations required to achieve complete saturation compromised patch stability (Fig. 2C). Fig. 2F shows the Schild analysis based on the obtained glutamate dose response curves (Fig. 2B-E). The solid lines in Fig. 2F are fits whose initial slopes were 0.99 (black) and 1.05 (red). As these were close to 1, they were constrained to unity to conform with the Schild equation. The intercept of these lines with the abscissa where (*EC*_50_ *_with NBǪX_ / EC*_50_ *_w/o NBǪX_* – 1) = 1 (indicated by the dashed horizontal line in Fig. 2F) provides *K_B_* values for NBǪX of 0.37 ± 0.02 μM for WT and 0.24 ± 0.07 μM for A622T mutant (*P* < 0.0001). The values are in close agreement with previously published K_NBǪX_ value of 0.59 μM for GluA2(Ǫ) WT, using a fluorescence-based Ca^2+^ imaging assay (Strange et al., 2006). The two NBǪX binding constants are not identical, but the difference between them is too small to account for ∼100-fold decrease in *EC*_50_ for glutamate for the mutant in the absence of NBǪX (*EC*_50_ for GluA2(Ǫ) WT 495 ± 43 μM and 5.4 ± 0.7 μM for GluA2(Ǫ) A622T, as measured from the steady-state responses). This suggests that *lc* in the GluA2(Ǫ) homomers primarily changes the efficacy with which the receptor undergoes conformational change(s) needed to open the ion channel rather than the binding affinity for glutamate. As *lc* mutation has so far been reported in patients in subunits GluA1 (de Ligt et al., 2012; Geisheker et al., 2017; Ismail et al., 2022; XiangWei et al., 2023) and GluA3 (Rinaldi et al., 2024), but not GluA2 (or GluA4), we next examined the effects of the equivalent mutation in GluA1 homomers.

**Figure 2.**
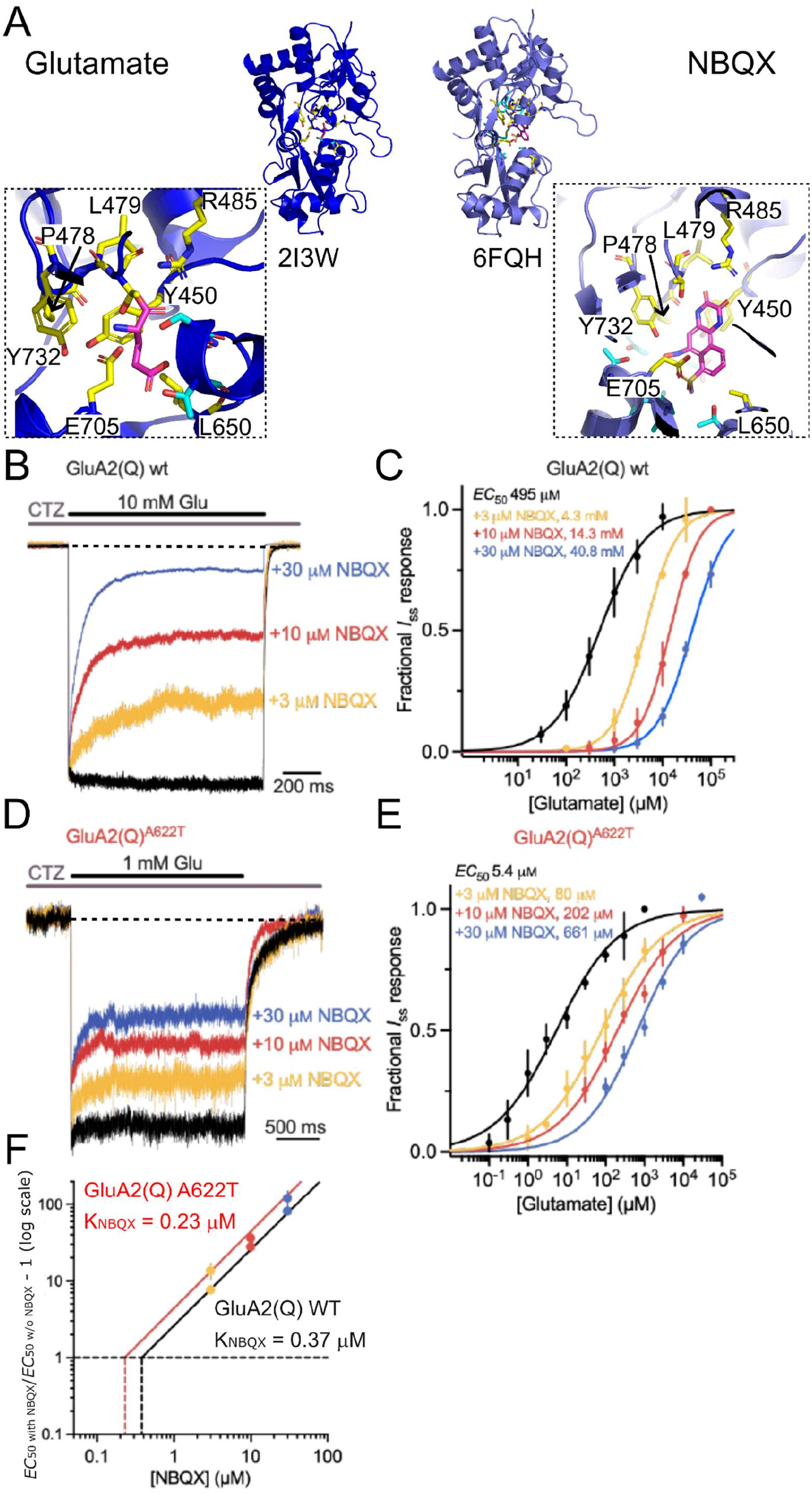
Schild analysis of wild-type and AC22T GluA2(Ǫ) AMPA receptors. (**A**) Glutamate and NBǪX binding site within GluA2 ligand-binding domain (LBD, shown in blue); Glu and NBǪX are in magenta, residues that interact with both, Glu and NBǪX (i.e. are within 4 Å of both ligands) are shown in yellow and their identity and position are indicated. Residues specific to only one interaction site are shown in cyan. Residue numbering same as in (Armstrong et al., 2006). PDB code for glutamate structure is 2I3W (Armstrong et al., 200C) and for NBǪX structure CFǪH (Coombs et al., 201S). **(B)** Outside-out patch clamp recordings from GluA2(Ǫ) WT receptors in HEK2S3 cells. Representative current traces evoked by glutamate (black bar, 10 mM, 1 s, –C0 mV) in the absence (black) or presence (coloured) of NBǪX (concentrations listed in the figure). 100 μM cyclothiazide (CTZ) is perfused throughout (grey bar). Amplitudes are standardised for illustrative purposes. **(C)** A series of dose-response curves for GluA2(Ǫ) WT receptors generated from I_ss_ responses to glutamate. Curves are generated from traces such as those presented in (B). The EC_50_ values were obtained from Hill-type fits and are provided alongside the concentrations they were obtained under. The glutamate EC₅₀ increased from 4S5 ± 43 µM in control (no NBǪX) to 4.3 ± 0.2, 14.3 ± 0.S, and 40.8 ± 1.7 mM in the presence of 3, 10 and 30 µM NBǪX, respectively. **(D)** Same experiment as in (A), but with GluA2(Ǫ) AC22T receptors (1 mM Glu, 2 s, –C0 mV). Note that longer glutamate pulses were used for GluA2(Ǫ) AC22T to accommodate the variant’s slower decay kinetics. **(E)** Dose-response curves for GluA2(Ǫ) AC22T mutant; examples of such traces are depicted in (D); n = 3 – C for each point. EC₅₀ increased from 5.4 ± 0.7 µM to 7S.S ± S.0, 202 ± 24.4, and CC1 ± 85.5 µM at the corresponding NBǪX concentrations of 3, 10 and 30 µM NBǪX, respectively. Error bars represent s.d.; some bars are smaller than the circle size. **(F)** Schild plot illustrating the action of NBǪX acting on GluA2(Ǫ) WT (black slope) and GluA2(Ǫ) AC22T mutant (red slope). Plots are produced from the data represented in (C) and (E). Lc mutation results in a 1.Cx decrease in K_NBǪX_ from 0.37 ± 0.02 μM (WT) to 0.24 ± 0.07 μM (AC22T) (n = 3-5 patches/concentration, extra sum-of-squares F-test, P < 0.0001). The data points (colour-coded as NBǪX concentrations in (C) and (E)) fall on a straight line whose slope is constrained to unity.

### *Lurcher* mutation decreases less profoundly desensitization in GluA1 homomers than GluA2(Ǫ) homomers

We next characterised *lc* mutation in GluA1 homotetramers to establish whether the effects of the mutation depend on the AMPA receptors subunit identity. Note that all GluA1 subunits throughout this study (and in vivo) are GluA1(Ǫ). Fig. 3 shows the same kinetic analysis of GluA1 *lc* homomers as was done for GluA2(Ǫ) *lc* homomers in Fig. 1. Similar to GluA2(Ǫ) A622T homomers, the GluA1 A636T mutation resulted in unchanged activation time, increased deactivation and desensitization time constants and decreased extent of desensitization when compared to GluA1 WT homomers. While the increase in the deactivation time constant was comparable to GluA2(Ǫ) *lc* homomers (*τ_deact_* = 39.9 ± 17.6 ms (n = 4) and 52.5 ± 31.0 ms (n = 13), respectively; *P* = 0.33), the changes in desensitization properties were less profound in GluA1 than GluA2(Ǫ) *lc* homomers (Fig. 3). GluA1 *lc* homomers desensitized with *τ_w, desens_* of 10.1 ± 4.0 ms (n = 11) and GluA2 *lc* homomers of 61.8 ± 47.2 ms (n = 15) (*P* < 0.001). The extent of desensitization was also larger in GluA1 than GluA2(Ǫ) *lc* homomers (*I_ss_/I_peak_* of 54.0 ± 13% (n = 12) and 90.0 ± 10% (n = 49), respectively, *P* < 0.0001) (Fig. 3B, E-F). Comparison of *lc* mutant kinetics in GluA1 and GluA2(Ǫ) homomers indicates that the mutation exerts similar, but not identical effects in the two subunit types, causing more profound changes in the desensitization of GluA2(Ǫ) homomers than GluA1 homomers (Table 1). The presence of the *lc* mutation appears to exacerbate already existing differences in the kinetics of WT GluA1 and GluA2(Ǫ) homomers (Table 1).

**Figure 3.**
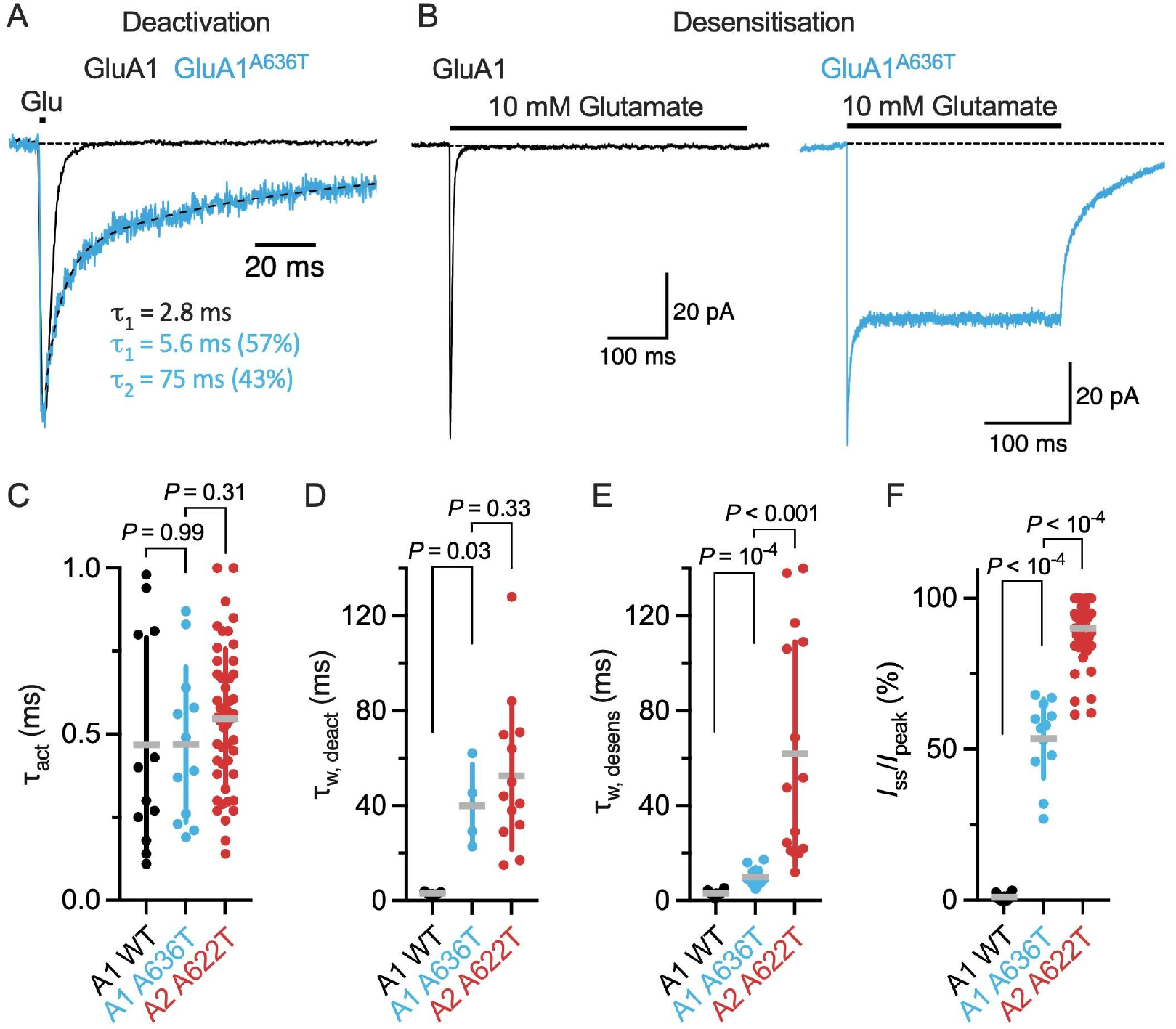
GluA1 wild-type (WT) and GluA1 AC3CT lurcher kinetics. (**A**) Representative traces from outside-out patch clamp recordings from GluA1 WT (black trace) and GluA1 AC3CT (blue trace) receptors in HEK-2S3 cells. The traces show deactivation currents generated from 1 ms applications of saturation glutamate (10 mM; –C0 mV; thick black bar). The exponential decays were fitted (dashed line) with one (WT) or two (AC3CT) components. The time constants and respective amplitudes from the fits are shown for the representative traces. Current amplitudes are scaled to the same value for illustrative purposes. **(B)** Desensitising current responses from ‘long’ glutamate exposure (400 ms and 200 ms for WT (black) and AC3CT (blue), respectively; 10 mM; –C0 mV). **(C)** Pooled τ_act_ data for GluA1 WT (black spheres) (n = 12), GluA1 AC3CT (blue spheres) (n = 12), and GluA2(Ǫ) AC22T (red spheres) (n = 48). **(D)** Pooled τ_w, deact_ data from GluA1 WT τ_w, deact_ = 2.S8 ± 0.8C ms(n = 5), GluA1 AC3CT τ_w, deact_ = 3S.S ± 17.C ms (n = 4) and GluA2(Ǫ) AC22T τ_w, deact_ = 52.5 ± 31.0 ms (n = 13) taken from 1 ms (or 2 ms in case of GluA2(Ǫ) lc) application of glutamate. A1 WT vs. A1 AC3CT: P = 0.03; A1 AC3CT vs. A2(Ǫ) AC22T: P = 0.33, two-tailed unpaired t-tests with Welch’s correction and P = 0.4C with randomisation test. **(E)** Pooled τ_w, des_ data for GluA1 WT τ_w, des_ = 3.1 ± 1.2 ms (n = 12), GluA1 AC3CT τ_w, des_ = 10.1 ± 4.0 ms (n = 11) and GluA2(Ǫ) AC22T τ_w, des_ = C1.8 ± 47.2 ms (n = 15) from 200 ms, 400 ms and 500 ms applications of glutamate, respectively. A1 WT vs. A1 AC3CT: P < 0.0001; A1 AC3CT vs. A2(Ǫ) AC22T: P < 0.001, two-tailed unpaired t-tests with Welch’s correction and P = 0.000S with randomisation test). **(F)** Pooled I_ss_/I_peak_ data for GluA1 WT I_ss_/I_peak_ = 1 ± 1% (n = 12), GluA1 AC3CT I_ss_/I_peak_ = 54.0 ± 13% (n = 12) and GluA2(Ǫ) AC22T I_ss_/I_peak_ = S0.0 ± 10% (n = 4S) from 200 ms, 400 ms and 500 ms applications of glutamate, respectively. A1 WT vs. A1 AC3CT: P < 0.0001; A1 AC3CT vs. A2(Ǫ) AC22T: P < 0.0001, Welch’s t-test and randomisation test. In (C-F), circles represent individual patches and horizontal and vertical lines indicate the mean ± s.d.

**Table 1.** Kinetics of wild-type (WT) and lurcher (lc) receptors for GluA2(Q) homomers, GluA1 homomers and GluA1/GluA2(R) heteromers.

|  | **τ<sub>act</sub> (ms)** | **τ<sub>deact</sub> (ms)** | **τ<sub>desens</sub> (ms)** | ***I*<sub>ss</sub> / *I*<sub>peak</sub> (%)** |
|---|---|---|---|---|
| **GluA1 WT** | 0.5 ± 0.3 (12*) | 3 ± 0.9 (5) | 3 ± 1 (12) | 1 ± 1 (12) |
| **GluA1 *lc*** | 0.5 ± 0.2 (12) | 40 ± 18 (4) | 10 ± 4 (11) | 54 ± 13 (12) |
| **GluA2(Q) WT** | 0.5 ± 0.2 (33) | 2 ± 1 (7) | 8 ± 3 (32) | 3 ± 2 (22) |
| **GluA2(Q) *lc*** | 0.5 ± 0.2 (48) | 52 ± 31 (13) | 62 ± 47 (15) | 90 ± 10 (49) |
| **GluA1<sup>*WT*</sup> / GluA2(R)<sup>*WT*</sup>** | 0.4 ± 0.2 (29) | 3 ± 1 (8) | 7 ± 3 (30) | 4 ± 4 (30) |
| **GluA1<sup>*lc*</sup> / GluA2(R)<sup>*WT*</sup>** | 0.5 ± 0.3 (14) | 7 ± 3.5 (6) | 18 ± 6 (13) | 33 ± 10 (13) |

\*Number of patches

To investigate the basis of this subunit-dependent difference in desensitisation, we next examined receptor activity at the single-channel level, asking how the disparity in macroscopic steady-state currents between GluA1 *lc* and GluA2(Ǫ) *lc* receptors might be reflected in their single-channel properties.

Outside-out patches containing single GluA2(Ǫ) WT, GluA2(Ǫ) *lc* or GluA1 *lc* receptor were excised from HEK293 cells. Representative currents recorded during applications of saturating glutamate (10 mM, 1.5 s, −80 mV) are shown in Fig. 4A-C. GluA2(Ǫ) WT receptors rapidly desensitised during sustained glutamate application, leaving only brief and intermittent open-state sojourns, spending most of the time in the closed, non-conducting states (Fig. 4A, n = 5). Comparable low-level activity has also been previously observed in single-channel recordings of GluA1 WT homomers (Banke et al., 2000). In contrast, both GluA1 and GluA2(Ǫ) *lc* receptors displayed sustained channel activity throughout glutamate application due to the desensitization block. To compare the open probability (*P_o_*) of the two *lc* mutants, we determined the amount of charge transferred during their single-channel activity as described in (Plested C Baranovic, 2016). Following complete dissociation of preincubated NBǪX, *P*_o_ was determined for GluA1 and GluA2(Ǫ) *lc* mutants from periods in which steady-state channel activity had been established. The amount of transferred charge for each mutant, was then compared to the theoretical maximum charge that would have been transferred if the receptor was fully open 100% of the time for which glutamate was bound. GluA2(Ǫ) *lc* receptors exhibited a *P*_o_ of 0.97 ± 0.02 (n = 9; Fig. 4B), whereas GluA1 *lc* receptors had a *P*_o_ of 0.82 ± 0.09 (n = 4; *P* = 0.0007, Fig. 4C-D). While detailed single-channel analysis is out of scope here, qualitative, visual inspection of the single-channel recordings clearly shows more frequent transitions to the closed and lower-conductance states in GluA1 *lc* compared to GluA2 *lc* (Fig. 4B-C). Single-channel recordings, thus, suggest, the smaller macroscopic steady-state current of GluA1 *lc* receptors might be, at least in part, attributed to their more frequent sojourns to closed and low-conductance states relative to GluA2(Ǫ) *lc* receptors.

**Figure 4.**
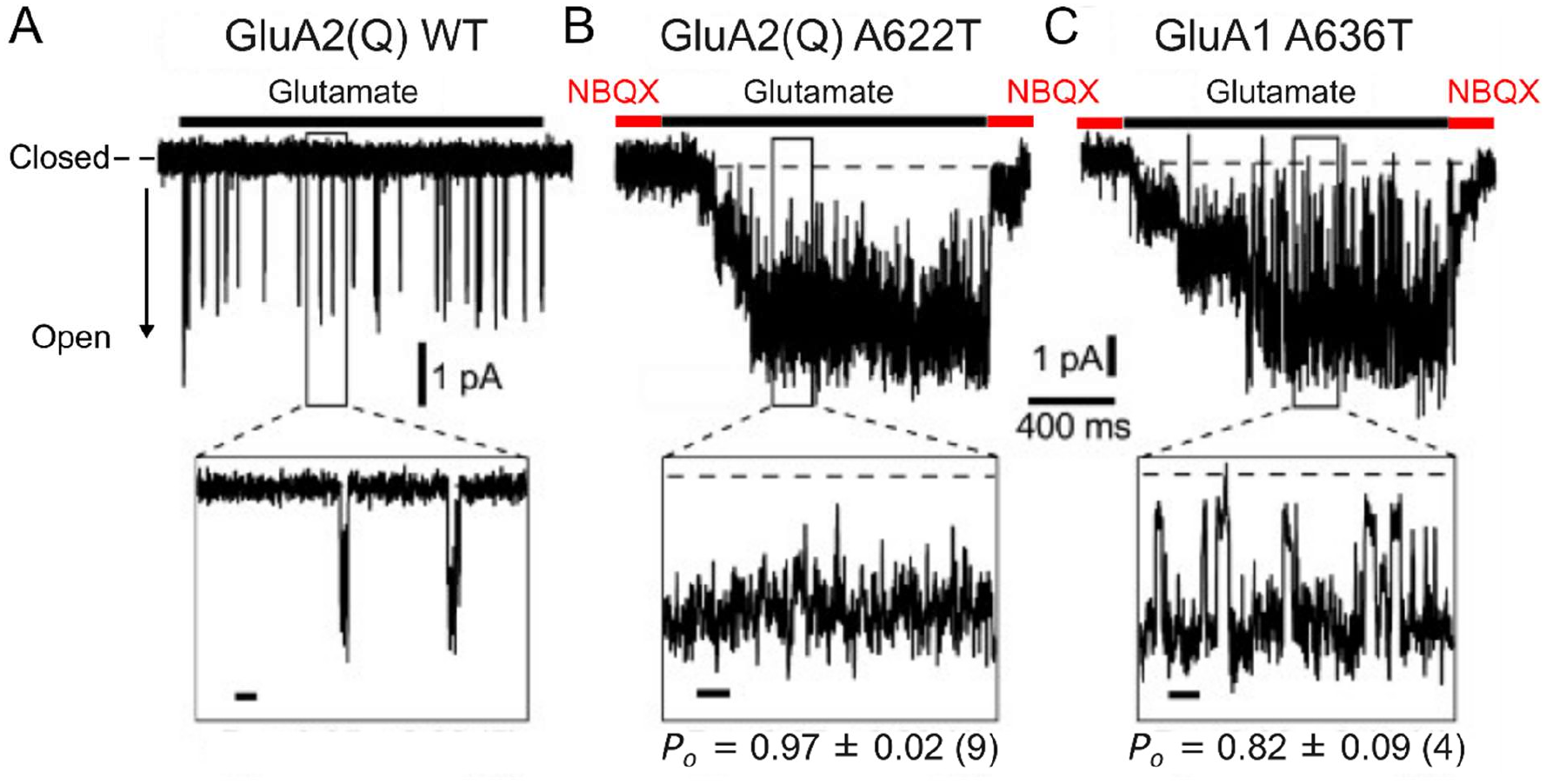
Effect of the lurcher mutation on single-channel open probability. (**A**) Representative current from an outside-out patch containing a single GluA2(Ǫ) WT receptor during application of glutamate (10 mM, 3 s, −80 mV). Insets show selected regions at an expanded timescale (scale bars, 20 ms). Recordings were sampled at 25 kHz and low-pass filtered at 1 kHz (Bessel filter). **(B)** Single-channel currents from a GluA2(Ǫ) AC22T receptor during glutamate application (10 mM, 1.5 s, −80 mV). Initial stepwise activation reflects dissociation of preincubated NBǪX (50 μM, 3 s) following exposure to glutamate. P_o_ was determined after complete NBǪX dissociation, once full channel activation had been established. **(C)** Same as B, but for GluA1 AC3CT. P_o_ values are shown in the respective panels (mean ± s.d.; n = number of patches). GluA2(Ǫ) AC22T vs. GluA1 AC3CT: P = 0.0007, unpaired t-test with Welch’s correction).

### Activating effects of the *lurcher* mutation are attenuated in GluA1*^lc^*/GluA2(R)*^WT^* heteromers compared to GluA1 *lc* and GluA2(Ǫ) *lc* homomers

In patients harbouring A636T mutation in the GluA1 subunit, the mutated subunit would be most prevalent in GluA1^A636T^/GluA2(R)^WT^ heteromers (GluA1*^lc^*/GluA2(R)*^WT^*) (Zhao et al., 2019). We have therefore, extended our analysis of the mutant to GluA1*^lc^*/GluA2(R)*^WT^* receptors. Before we could determine kinetics of GluA1*^lc^*/GluA2(R)*^WT^* receptors, we had to ensure that GluA2(R) WT and GluA1 *lc* subunits were indeed assembling as GluA1*^l^*^c^/GluA2(R)*^WT^* heteromers and not GluA2(R) WT or GluA1 *lc* homomers in our patches. GluA2(R) WT homomers have extremely low conductance (few hundred fS (Swanson et al., 1997)) and can therefore, even if assembled, be excluded as contributors to the current. The same is true for GluA2(R) *lc* homomers – we did not manage to get any currents from HEK cells transfected with GluA2(R) *lc* homomers. GluA1 *lc* homomers produce sizable currents (Fig. 3), but because they do not contain Arg residues in the pore, their currents can be blocked by polyamine spermine in a voltage-dependent manner. With 50 µM spermine in the pipette solution and at positive voltages (+60 mV or below), GluA1 *lc* homomers (and GluA1 WT homomers in controls) will be blocked by spermine and hence any visible currents in these conditions, will be from GluA1*^l^*^c^/GluA2(R)*^WT^* heteromers (or GluA1*^WT^*/GluA2(R)*^WT^* in controls) (Fig. 5A-B). For this reason, all the current traces for the kinetic analysis of GluA1*^lc^*/GluA2(R)*^WT^* heteromers were obtained in the presence of 50 µM spermine and at +60 mV (Fig. 5C-D). Just like GluA1 *lc* homomers and GluA2(Ǫ) *lc* homomers, GluA1*^lc^*/GluA2(R)*^WT^* heteromers also do not show any change in the activation time constant compared to the WT control (GluA1*^WT^*/GluA2(R)*^WT^*) (Fig. 5E and Table 1) (τ_w_,_act_ of 0.44 ± 0.20 ms (n = 29) and 0.53 ± 0.30 ms (n = 14), for wild-type and GluA1*^lc^*/GluA2(R)*^WT^* receptors, respectively (*P* = 0.30). Deactivation takes marginally longer in GluA1*^lc^*/GluA2(R)*^WT^* compared to GluA1*^WT^*/GluA2(R)*^WT^* heteromers (Fig. 5F and Table 1) (τ_w, deact_ of 6.97 ± 3.47 ms (n = 6) for GluA1*^lc^*/GluA2(R)*^WT^*, compared with 3.11 ± 1.19 ms for WT heteromers (n = 8, *P* = 0.041). Just like for homomeric receptors, the biggest changes are in the time course and extent of desensitization. The presence of *lc* mutation in GluA1 subunit of GluA1*^lc^*/GluA2(R)*^WT^* heteromers slows down desensitization and decreases the extent of desensitization compared to WT heteromers (Fig. 5G-H and Table 1) (τ_w, desens_ of 17.6 ± 5.90 ms (n = 13) for GluA1*^lc^*/GluA2(R)*^WT^*, compared with 6.70 ± 3.12 ms for wild-type heteromers (n = 30; *P* < 0.0001). The fractional steady-state current (I_ss_/I_peak_) likewise increased from 4 ± 4% to 33 ± 10% (*P* < 0.0001). Both effects are less profound in GluA1*^lc^*/GluA2(R)*^WT^* heteromers than in GluA1 *lc* or GluA2(Ǫ) *lc* homomers (Table 1).

**Figure 5.**
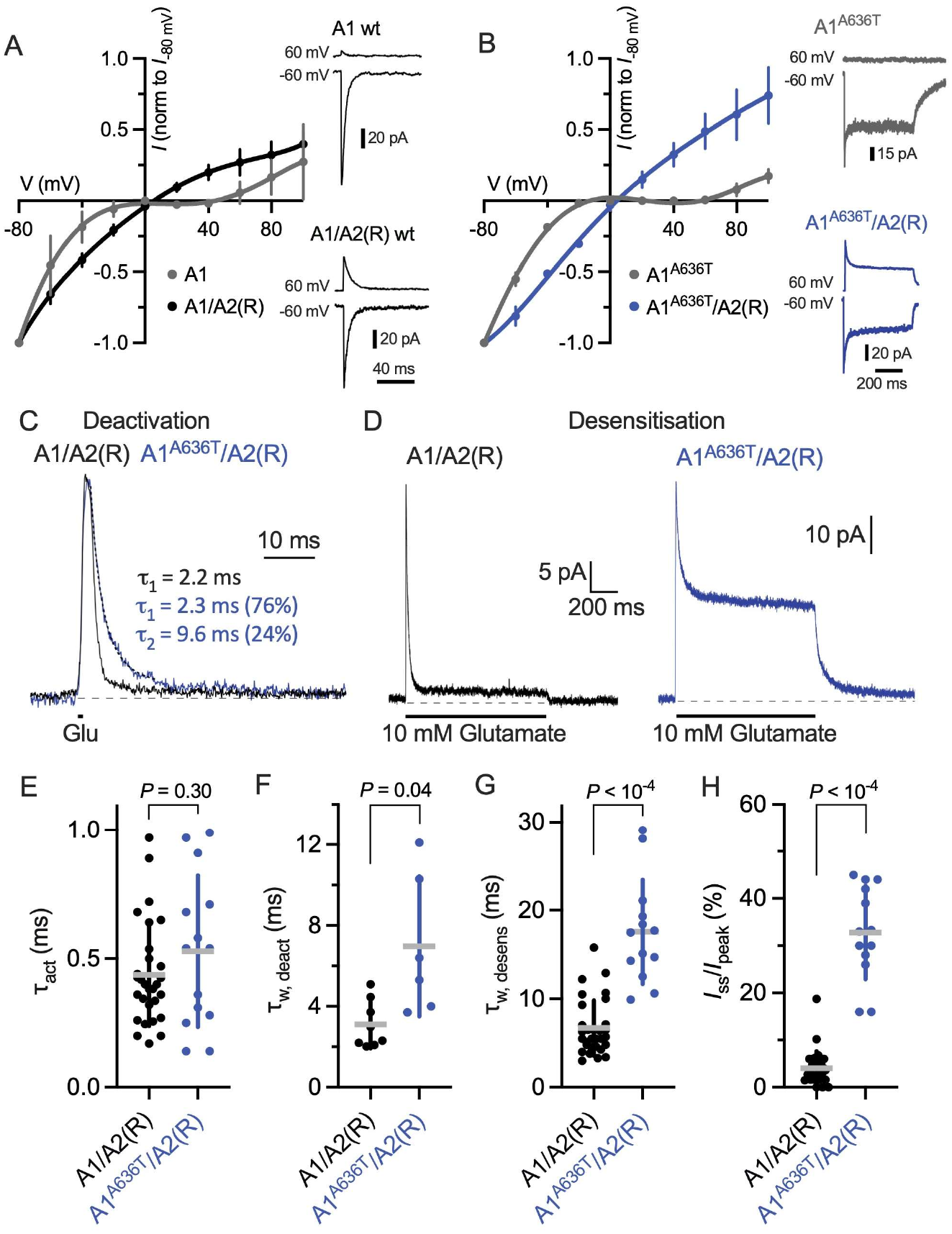
Effects of lurcher mutation are attenuated in GluA1 AC3CT/GluA2(R) WT heteromers. (**A**) Current-voltage relationships for WT GluA1 homomers and GluA1/GluA2(R) heteromers recorded in the presence of intracellular spermine (50 µM). Currents evoked by saturating glutamate (10 mM) over membrane potentials from –80 to +100 mV were normalised to the current at –80 mV. Representative currents recorded at –C0 and +C0 mV are shown alongside. **(B)** Same as (A), but for GluA1^AC3CT^ homomers and GluA1^AC3CT^/GluA2(R)^WT^ heteromers. The reduced inward rectification of GluA2(R)-containing receptors in (A) and (B) was used to distinguish heteromeric from GluA1 homomeric currents. **(C)** Representative deactivation currents from GluA1/GluA2(R) wild-type and GluA1^AC3CT^/GluA2(R) receptors following brief (2 ms) application of saturating glutamate (10 mM, +C0 mV). Decays were fitted with one or two exponentials; time constants (τ_1_, τ_2_) and their relative weights (parentheses) for the representative traces are indicated. **(D)** Representative desensitising currents from GluA1/GluA2(R) WT and GluA1^AC3CT^/GluA2(R) receptors during prolonged application of saturating glutamate (10 mM, +C0 mV). **(E)** Pooled weighted activation data: WT heteromers τ_w_, _act_ = 0.44 ± 0.20 ms (n = 2S), GluA1^lc^/GluA2(R)^WT^ τ_w_, _act_ = 0.53 ± 0.30 ms (n = 14), P = 0.30, Welch’s t-test and P = 0.23 with randomisation test. **(F)** Pooled weighted deactivation data: WT heteromers τ_w_, _deact_ = 3.11 ± 1.1S ms (n = 8), GluA1^lc^/GluA2(R)^WT^ τ_w_, _deact_ = C.S7 ± 3.47 ms (n = C), P = 0.041, Welch’s t-test and P = 0.005 with randomisation test. **(G)** Pooled weighted desensitization data: WT heteromers τ_w_, _desens_ = C.70 ± 3.12 ms (n = 30), GluA1^lc^/GluA2(R)^WT^ τ_w_, _deact_ = 17.C ± 5.S0 ms (n = 13), P < 0.0001, Welch’s t-test and randomisation test. **(H)** Pooled fractional steady-state current data: WT heteromers τ_w_, _desens_ = 4 ± 4% (n = 30), GluA1^lc^/GluA2(R)^WT^ τ_w_, _deact_ = 33 ± 10% (n = 13), P < 0.0001, Welch’s t-test and randomisation test. Circles represent individual patches and horizontal and vertical lines indicate mean ± s.d.

Due to the prevalence of GluA1*^lc^*/GluA2(R)*^WT^* (Geisheker et al., 2017; Zhao et al., 2019), we have also measured the rate of recovery from desensitization and *EC*_50_ for glutamate (Figure 6). Recovery from desensitization was faster in GluA1*^lc^*/GluA2(R)*^WT^* heteromers compared to GluA1*^WT^*/GluA2(R)*^WT^* (τ_rec_ = 31.5 ± 1.8 ms, n = 3, versus 62.4 ± 3.8 ms, n = 3, *P* < 0.0001) (Fig. 6A-B), while *EC*_50_ value was smaller in GluA1*^lc^*/GluA2(R)*^WT^* compared to the WT controls (57.1 ± 23.7 µM, n = 3, versus 551 ± 190 µM, n = 4, respectively; *P* = 0.001) (Fig. 6C-D). Both results are consistent with *lc* mutation promoting AMPA receptor activity. Increased sensitivity to glutamate in GluA1*^lc^*/GluA2(R)*^WT^* (i.e. decrease in *EC*_50_ by an order of magnitude) is consistent with the effects of *lc* in GluA1 (Klein C Howe, 2004) and in GluA2(Ǫ) homomers (decrease in *EC*_50_ by three orders of magnitude for peak responses (Fig. 1H-I) and two orders of magnitude for steady-state responses (Fig. 2D-E)). Faster recovery from desensitization agrees with the general *lc* effect of reducing desensitization in AMPA receptors via slower time course of desensitization and decreased extent of desensitisation in both, homomers (Figs. 1C, F-G and 3B, E-F) and heteromers (Fig. 5D, G-H).

**Figure 6.**
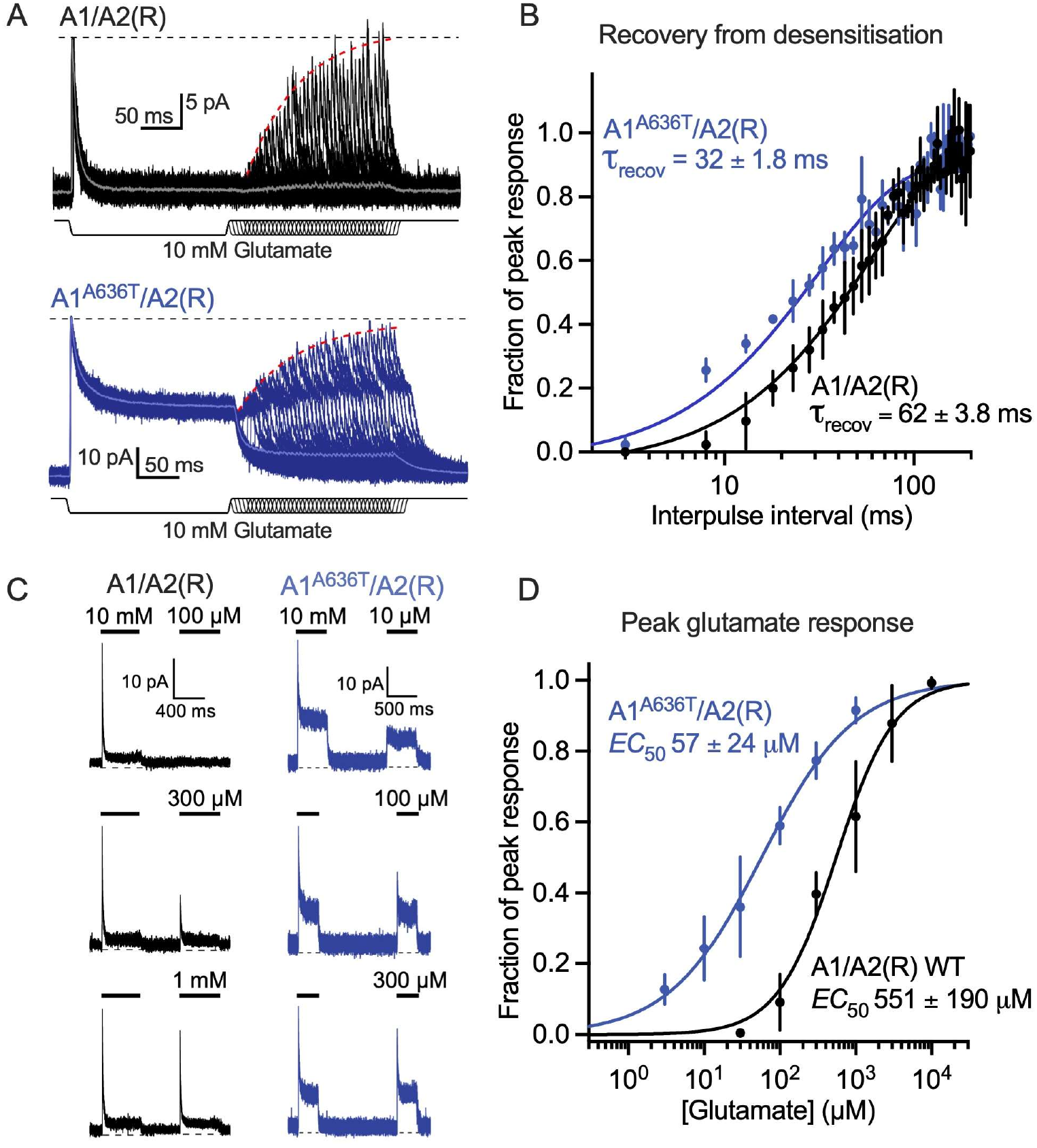
GluA1AC3CT/GluA2(R)^WT^ recovers faster from desensitisation and has lower EC50 for glutamate than WT heteromers. (**A**) Representative paired-pulse recordings from GluA1/GluA2(R) WT (black) and GluA1^AC3CT^/GluA2(R)^WT^ receptors (blue), recorded at +C0 mV in the presence of intracellular spermine (50 µM). A conditioning application of glutamate (10 mM, 200 ms) was followed by a 10 ms test pulse at increasing interpulse intervals. Lighter traces represent averaged responses, and dashed red lines indicate fits to the test responses. **(B)** Pooled recovery from desensitisation for GluA1/GluA2(R) WT (black) and GluA1^AC3CT^/GluA2(R)^WT^ receptors (blue), obtained from peak responses such as those shown in (A). Curves were fitted with a Hodgkin–Huxley-type recovery function, yielding recovery time constants (τ_recov_) of C2.4 ± 3.8 ms (n = 3, S5% CI, 4S.8-C8.1 ms) and 31.5 ± 1.8 ms (n = 3, S5% CI, 28.1-40.3 ms), respectively (P < 0.0001, extra sum-of-squares F test). **(C)** Representative macroscopic currents from GluA1/GluA2(R) WT and GluA1^AC3CT^/GluA2(R)^WT^ receptors evoked over a range of increasing glutamate concentrations. **(D)** Glutamate concentration-responses constructed from peak currents, obtained from recordings such as those shown in (C), for GluA1/GluA2(R) WT (black) and GluA1^AC3CT^/GluA2(R)^WT^ receptors (blue). Hill-type fits yielded EC₅₀ values of 551 ± 1S0 µM (n = 4) and 57.1 ± 23.7 µM (n = 3), respectively (P = 0.001, unpaired t-test with Welch’s correction). Error bars represent s.d.; where not visible, they are smaller than the symbols. τ_recov_ values are reported as best-fit estimates ± s.e.

### *IC_50_* for perampanel in *lurcher* mutants depends on subunit type and the presence of auxiliary subunit stargazin

We next determined how *lc* mutation affects *IC_50_* of perampanel (PM)P), the only commercially available drug targeting AMPA receptors. PMP is a non-competitive antagonist of AMPA receptors with 4 molecules of PMP binding to the transmembrane helices, just above the extracellular membrane leaflet per one receptor (Fig. 7A-C). The binding site for PMP is dominated by one subunit with only one residue contributing from the neighbouring subunit (Yelshanskaya et al., 2016) (Fig. 7B). Although the binding site involves residues from the SYTANLAAF motif (Tyr616, Leu620 and Phe623), the *lc* residue (Ala622) does not seem to directly contribute to PMP binding (Fig. 7B-C), based on the 5L1F structure (Yelshanskaya et al., 2016) which has a limited resolution of ∼4 Å. However, due to its proximity to the PMP binding site, we thought *lc* mutation might affect *IC*_50_ of PMP (i.e. PMP concentration which inhibits 50% of the maximal AMPA receptor response). Indeed, PMP *IC*_50_ (measured from the peak responses) for GluA2(Ǫ) WT receptors is 0.49 ± 0.03 μM (n = 3-7 patches per concentration) and shifts to 52.4 ± 6.2 µM in GluA2(Ǫ) *lc* receptors (n = 3-5) (Fig. 7D-E), corresponding to an approximately 100-fold reduction (*P* < 0.0001). The right-shift is expected given the *lc* propensity to keep AMPA receptors active. The shift is more modest in GluA1*^lc^*/GluA2(R)*^WT^* heteromers where PMP *IC*_50_ for WT is 0.51 ± 0.04 μM (n = 3) and for mutant 1.27 ± 0.15 μM (n = 3-4), corresponding to an approximately 2.5-fold shift (*P* < 0.0001) (Fig. 7F-G). The difference decreases even further if the GluA1/GluA2(R) heteromers are associated with auxiliary subunit γ-2 (from 0.20 ± 0.02 µM for WT (n = 3-4) to 0.46 ± 0.03 μM for the mutant (n = 3-5)), representing an approximately 2.3-fold decrease (Fig. 7H-I). As above (Figs. 5 and 6), heteromer recordings were obtained in the presence of 50 μM spermine in the pipette solution and at positive voltages to isolate heteromer currents. TARP γ –2, however, partially alleviates spermine block of calcium-permeable AMPA receptors (Carbone C Plested, 2016; Soto et al., 2007), which means some of the currents in heteromer recordings with y-2 (Fig. 7H) could be contributed by GluA1 homomers associated with y-2.

**Figure 7.**
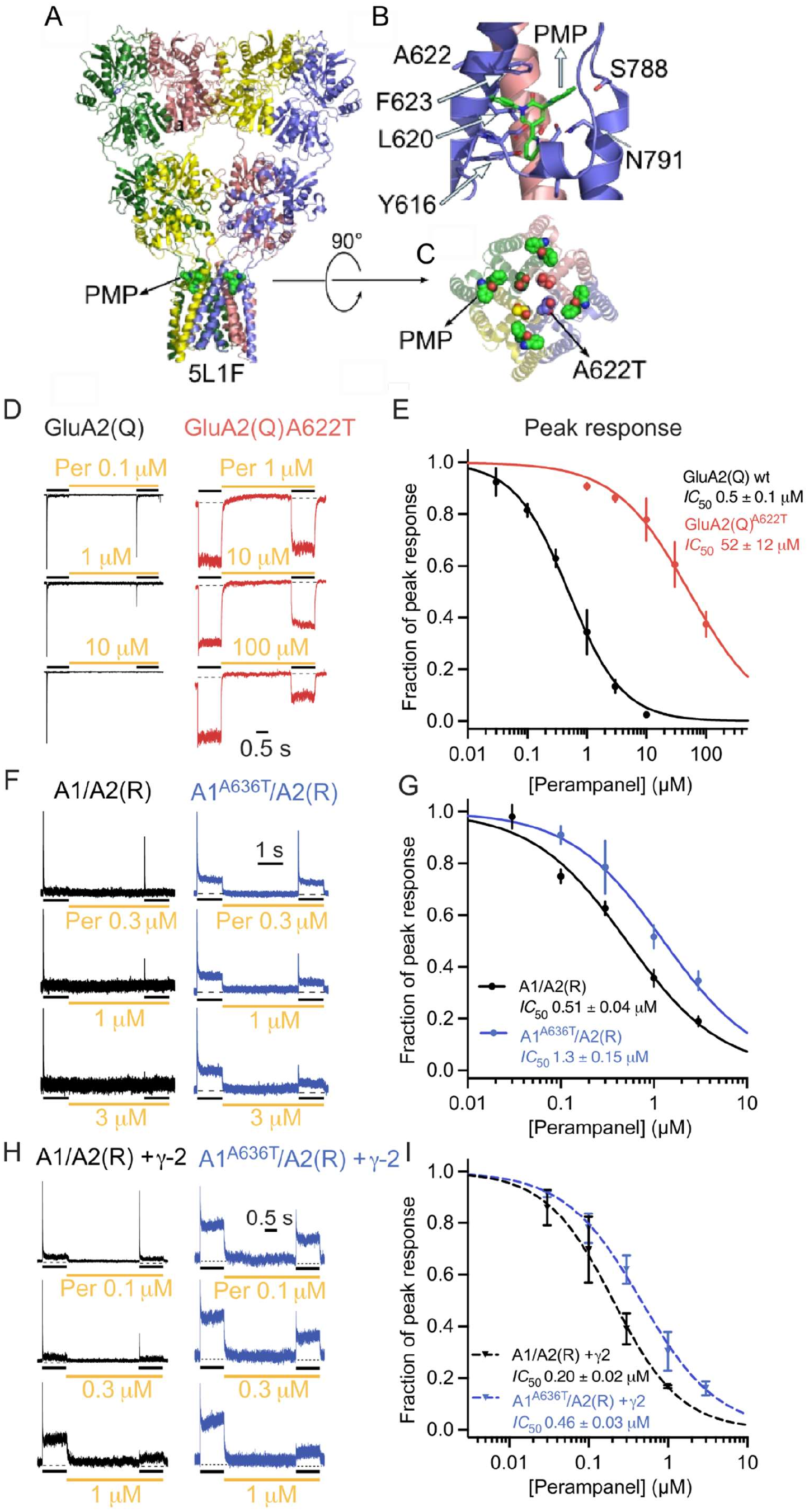
Perampanel IC50 and lurcher mutations in GluA2(Ǫ) homomers and GluA1^lc^/GluA2(R)^WT^ heteromers with and without auxiliary subunit y-2. (**A**) Structure of GluA2(Ǫ) homomer with individual subunits colour-coded (in green, red, blue and yellow) and four bound perampanel (PMP) molecules shown as green spheres (PDB: 5L1F (Yelshanskaya, Neuron, 201C)). **(B)** Zoom into the binding site of one PMP molecule (green sticks) formed by subunits B and C (red and blue). Interacting residues are indicated and their side-chains shown as sticks. Lc residue (AC22) is indicated although it does not form the binding site. **(C)** Top-down view of the PMP (green spheres) binding sites with lc residue in each subunit shown as spheres. GluA2 extracellular domains are removed for clarity, and only transmembrane region is shown. In all sphere representations, oxygen atoms are shown in red and nitrogen in blue. **(D)** Representative glutamate-evoked currents from GluA2(Ǫ) WT (black) and GluA2(Ǫ) AC22T receptors (red) following preincubation (4 s) with increasing concentrations of PMP, as indicated. Peak (conditioning) responses were evoked by saturating glutamate (10 mM for WT and 1 mM for GluA2(Ǫ) AC22T; 500 ms, −C0 mV). **(E)** PMP dose-response curves constructed from the fractional inhibition of peak current responses obtained from recordings such as those shown in (D). Hill-type fits yielded IC₅₀ values of 0.4S ± 0.03 µM and 52.4 ± C.2 µM for GluA2(Ǫ) WT and GluA2(Ǫ) AC22T receptors, respectively (n = 3-7 and 3-5; P < 0.0001, extra sum-of-squares F test). **(F)** Representative traces showing glutamate-responses (10 mM, +C0 mV) from GluA1/GluA2(R) WT (black) and GluA1 AC3CT/GluA2(R) receptors (blue) following preincubation with the indicated concentrations of PMP. **(G)** PMP dose-response curves for GluA1/GluA2(R) WT (black) and GluA1^AC3CT^/GluA2(R)^WT^ receptors (blue). Hill-type fits yielded IC₅₀ values of 0.51 ± 0.04 µM (n = 3) and 1.27 ± 0.15 µM (n = 3-4) for WT and lc heteromers, respectively. **(H-I)** As in (F-G), but in the presence of auxiliary subunit γ-2. IC_50_ values were 0.20 ± 0.02 µM (n = 3-4) for WT and 0.4C ± 0.03 µM (n = 3-5) for lc heteromers. The mutation produced shifts in PMP IC₅₀ both in the absence and presence of γ-2 (P < 0.0001, extra sum-of-squares F tests). Symbols represent mean fractional peak responses, and error bars denote s.d.; where not visible, error bars are smaller than the symbols. IC₅₀ values are reported as best-fit estimates ± s.e.

## Discussion

Since its first report in delta ionotropic glutamate receptors (δ2 iGluRs) (Kohda et al., 2000; Wollmuth et al., 2000), *lc* mutation has been characterised in various subunits in NMDA (Hu C Zheng, 2005a, 2005b; Murthy et al., 2012), kainate (Schwarz et al., 2001) and AMPA receptors (Ismail et al., 2022; Klein C Howe, 2004; Rinaldi et al., 2024; Schwarz et al., 2001; Taverna et al., 2000; Zhang et al., 2017). In most of these cases and certainly in all AMPA receptor subunits, *lc* leads to an overactive receptor, but beyond that, our understanding of its mechanism and effects on native AMPA receptor complexes remain incomplete. The relevance of the *lc* mutation in AMPA receptors has recently been renewed as it was found in 2 out of 4 AMPA receptor genes (*GRIA1* and *GRIA3*) in patients diagnosed with neurodevelopmental disorders (de Ligt et al., 2012; Geisheker et al., 2017; Ismail et al., 2022; Rinaldi et al., 2024; XiangWei et al., 2023; Xu et al., 2026). Here, we show that *lurcher’s* activating effects arise primarily from altered gating efficacy, and that its impact on receptor kinetics and pharmacology is reduced in more physiological complexes containing heteromeric AMPA receptors and auxiliary subunit γ-2.

Our analysis of kinetics in GluA2(Ǫ) *lc* (A622T) homomers shows no change in the activation, about 50-fold slower deactivation, about 7-fold slower desensitization and almost complete block of desensitization compared to GluA2(Ǫ) WT homomers. These results are consistent with the effects of the *lc* mutation in GluA1 ((Chen et al., 2025; Ismail et al., 2022; Klein C Howe, 2004; Taverna et al., 2000) and Fig. 3), GluA3 (Rinaldi et al., 2024) and GluA4 homomers (Zhang et al., 2017). Although the overall effects of *lc* are comparable between different subunits, the extent of change does vary from one subunit type to the other. We show that *lc* in GluA1 subunit results in about 50% block of desensitization compared to GluA2 homomers where the block is nearly 100% (Table 1). Both, deactivation and desensitization time courses are slowed down less profoundly in GluA1 *lc* homomers than in GluA2 *lc* homomers compared to their WT counterparts (deactivation is about 13 times slower in GluA1 *lc* homomers and about 26 times in GluA2 homomers whereas desensitization is about 3-fold slower in GluA1 and 8-fold in GluA2 homomers) (Table 1). These differences are expected given that kinetics of AMPA receptor homomers will vary depending on the subunit type, with GluA2 homomers, for example, exhibiting the slowest desensitization (Klein C Howe, 2004; Rinaldi et al., 2024; Zhang et al., 2017). In fact, our measurements (Table 1) and a previous report (Klein C Howe, 2004) show that GluA1 WT homomers appear to deactivate and desensitize at similar rates.

Unsurprisingly, the presence of *lc* in only two subunits in GluA1*^lc^*/GluA2(R)*^WT^* heterodimers further attenuates *lc* effects, as has been shown previously for GluA1 flip(R)*^WT^*/GluA2 flop(Ǫ)*^lc^* (Schwarz et al., 2001) and GluA2 flip(R)*^WT^*/GluA3 flip(Ǫ)*^lc^* (Rinaldi et al., 2024) heteromers. Clearly, smaller kinetic effects are expected in GluA1*^lc^*/GluA2(R)*^WT^* heterodimers compared to homomers, as only 2 subunits carry the mutation, but two additional factors might contribute to this attenuation. Firstly, within a tetramer, GluA1 subunits occupy positions closer to the overall 2-fold axis of symmetry at the level of ligand-binding domains (Herguedas et al., 2019; Zhao et al., 2019). Structural models indicate subunits at these positions contribute less to channel opening than those further away from the central axis (Twomey et al., 2017). Secondly, the Arg residue in the selectivity filter of GluA2 subunits in the GluA1*^lc^*/GluA2(R)*^WT^* heteromer will further reduce overall conductance (Heiser et al., 2025).

The presence of *lc* in GluA1 subunit, in GluA1*^lc^*/GluA2(R)*^WT^* heteromers associated with auxiliary subunit γ-2 has a relatively mild effect on *IC*_50_ of PMP, increasing its value about 2 times compared to the WT complexes. The modest shift in PMP *IC*_50_ in heteromers associated with γ-2 suggests PMP could be as effective in patients carrying the GluA1 *lc* mutation as in those with WT receptors, although the presence of other auxiliary subunits than y-2 will shape sensitivity to PMP. CNIH2, and to a lesser extent γ-5, have both been shown to reduce PMP potency in GluA2-containing complexes (Gangwar et al., 2023). Majority of GluA1/GluA2 WT heteromers in rat brain appear to be associated TARP γ-8 and CNIH2 (Yu et al., 2021; Zhao et al., 2019) and how *lc* affects these complexes remains to be determined.

Given the clinical importance of the *lc* mutation, we set out to determine whether it changes the binding affinity for glutamate or the efficacy with which glutamate opens the ion channel. Our Schild analysis reveals about 1.5-fold decrease in the binding constant for NBǪX, a competitive AMPA receptor antagonist, for GluA2(Ǫ) *lc* homomers compared to WT. This cannot explain the large decrease in *EC*_50_ for glutamate caused by the *lc* mutation in GluA2(Ǫ) homomers (by three orders of magnitude for peak responses and two orders of magnitude for steady-state responses). The shallowing of the Hill slope in the glutamate dose-response curve of the *lurcher* mutant further supports change in gating efficacy (Colquhoun, 1998). We therefore conclude that *lc* mutation primarily increases the efficacy with which AMPA receptors open, once glutamate is bound. From the structural perspective, this is not surprising as *lc* mutation is near the ion channel gate and, based on the structural models, appears to influence conformation of the gate more than of the remote ligand-binding domains (L. Chen et al., 2014; Kumar Mondal et al., 2025). Previous attempt to explain the mechanism behind *lc* concluded it decreased dissociation rate of glutamate from the ligand-binding domains based on the modelling of macroscopic currents (Klein C Howe, 2004). This might seem to be in disagreement with our data, however, the authors note that what appears in their model as a large decrease in the rate of glutamate unbinding, “*is, in fact, the effect of the mutation to stabilize a state that is inherently unstable in wild-type channels, namely the closed-bound transition state that precedes channel gating*” (Klein C Howe, 2004), thus suggesting their model does not exclude the possibility that *lc* primarily affects efficacy of channel opening. Although Schild analysis offers an elegant way of determining the microscopic binding constant of competitive antagonists, it is not without pitfalls. Mostly, it relies on the antagonist being a true competitive antagonist (i.e. that it binds with same affinity to the resting and active conformations of the receptor) rather than an inverse agonist (which binds with higher affinity to the resting state than open state conformations). In most cases, this is inconsequential unless the receptor can be constitutively active, i.e. open without agonist binding (Colquhoun, 1973). Whether *lc* can open without ligand binding remains unclear: δ2 iGluRs carrying *lc* mutation have been characterized as constitutively active in (Kohda et al., 2000; Zuo et al., 1997), however, (Klein C Howe, 2004) suggest that *lc* does not make AMPA receptors constitutively active, but instead, makes them more sensitive to ambient glutamate. The best way to resolve this would be to determine microscopic binding and gating constants from single-channel recordings.

Our study focuses on a change of the third Ala into Thr within the SYTANLAAF motif as this was a hot spot identified in patients with various degrees of neurodevelopmental delay (Geisheker et al., 2017; XiangWei et al., 2023), however, other mutations have been reported in patients within the SYTANLAAF motif across AMPA receptor subunits (Coombs et al., 2022; Davies et al., 2017; Martin et al., 2017; Rinaldi et al., 2024). AMPA receptor SYTANLAAF motif, thus, appears regularly mutated in patients, with most, but not all mutations resulting in an increase in overall AMPA receptor activity. In vivo, other factors, such as receptor expression, assembly and trafficking will combine with the kinetics to shape the phenotype of a mutation (C.-M. Chen et al., 2025; Kim et al., 2010; XiangWei et al., 2023; Xu et al., 2026). Nevertheless, given the growing numbers of patients with mutations in ionotropic glutamate receptor genes and the lack of treatments (Meagan Collins et al., 2021), our data showing the subunit– and auxiliary-subunit dependence of the *lc* mutant as well as its modulation by perampanel advances our understanding of this disease-associated mutant.

## Methods

### Heterologous Expression

Plasmids for the GluA1(Ǫ) flip WT, GluA2(Ǫ) flip WT, GluA2(R) flip WT and GluA2(Ǫ) flip A622T genes were on the pRK5 background with eGFP gene downstream of an internal ribosomal entry site sequence (IRES). All constructs corresponded to the flip splice isoform, which confers slower desensitisation kinetics than the corresponding flop variant (Koike et al., 2000), and contained the edited R/G site. For GluA2(R) construct, the fluorescent reporter was switched from eGFP to mCherry. Plasmid carrying mouse γ-2 gene was on the pRK8 background containing the non-cytotoxic version of dsRed also under IRES control (dsRed-Max, Addgene catalog number 21718) (Strack et al., 2008). All 5 plasmids were a kind gift from Andrew Plested (Humboldt University, Berlin, Germany). *Lurcher* mutation was introduced into the GluA2(R) flip WT and GluA1(Ǫ) flip (A622T and A636T respectively) using ǪuikChange Lightning Site-Directed Mutagenesis kit (Agilent Technologies) and confirmed by sequencing.

Recombinant wild-type and *lc* mutant AMPA receptors were expressed in Human Embryonic Kidney (HEK)-293 cells (Sigma). The cells were cultured in Minimum Essential Medium (MEM) (Gibco) supplemented with 10% (v/v) Fetal Bovine Serum (FBS) (Gibco), 2 mM L-glutamine (Gibco), and 50 U/mL penicillin-streptomycin (Gibco), following standard protocols in a 5% CO₂ incubator at 37°C (Plested C Baranovic, 2016). For electrophysiological recordings, cells were seeded at a density of 3 × 10⁶ on 13 mm borosilicate glass coverslips (Avantor), pre-coated with poly-L-lysine (4.5 µg/mL) (Sigma-Aldrich). The next day, transient transfection of plasmid DNA constructs was performed using either calcium phosphate method or, in case of single-channel recordings and constructs with low expression efficiency, Lipofectamine 3000 (Invitrogen). GluA1/GluA2 and γ-2 were transfected at a mass ratio of 1:1:2. Following transfection, cells were treated with the competitive antagonist, 2,3-dioxo-6-nitro-7-sulfamoyl-benzo[f]quinoxaline (NBǪX) (Tocris Bioscience) at 50 μM to reduce excitotoxicity. Electrophysiological recordings were conducted 18 to 74 hours post-transfection.

### Electrophysiology

Coverslips coated with cells were transferred to an inverted microscope (IX73 Olympus), and eGFP and/or mCherry-positive cells were selected for patch-clamp recordings using a *p*E-300^lite^ LED light source. Recordings were made in the following external solution (mM): 150 NaCl, 5 HEPES, 0.1 MgCl_2_, 0.1 CaCl_2_, pH 7.3, 280 mOsm. For macroscopic recordings, patch-clamp electrodes fabricated from borosilicate glass (1.5 mm outer diameter (o.d.), 1.17 mm inner diameter (i.d.) (Harvard Apparatus), were fire-polished to a final resistance of 3-8 MΩ. Internal pipette solutions contained (mM): 115 NaCl, 10 NaF, 5 Na_4_BAPTA, 0.5 CaCl_2_, 1 MgCl_2_, 5 HEPES, and 10 Na_2_ATP, pH 7.3 with NaOH, 280-300 mOsm. Recordings were made at 22-25°C from outside-out patches using an Axopatch 200B amplifier (Molecular Devices). Signals were low pass filtered at 5 kHz, digitised at 20 kHz using a Dendrite data acquisition system (Sutter Instruments), and analysed using SutterPatch software (version 3.2.1.) (Sutter Instruments) unless stated otherwise. Currents were recorded at –60 or –40 mV unless specified otherwise.

Single-channel recordings were performed as described in (Plested C Baranovic, 2016). Recording pipettes had resistance of 10-20 MΩ and were coated with Sylgard. Outside-out patches were voltage-clamped at −80 mV. For *lc* receptors, the patches were first incubated in NBǪX (50 µM) for 3 s, before rapidly switching to a glutamate-containing solution (10 mM) without NBǪX. The incubation in NBǪX made it easier to find a stable baseline given the high activity of *lc* mutants. GluA2 WT receptors were not incubated in NBǪX.

For macroscopic current recordings, patches were discarded if the baseline leak current exceeded 10% of the peak agonist-evoked current. For single-channel recordings, patch quality was monitored via the root-mean-square current noise (I_RMS_) indicated on the amplifier. Prior to a single-channel recording, a baseline I_RMS_ was measured with the electrode positioned immediately adjacent to the bath solution; recordings were only initiated if this baseline noise was below 200 fA. During data acquisition, recordings were aborted and discarded if the I_RMS_ exceeded 500 fA.

### Rapid Ligand Application to Excised Patches

Sub-millisecond ligand application was achieved by rapidly alternating between continuously flowing solutions. Solution exchange was controlled by the lateral movement of a custom-built four-barrel application tool, fabricated from borosilicate glass (CM Scientific) (Plested C Poulsen, 2021) and mounted on a piezoelectric translator (Physik Instrumente). The speed of solution exchange was assessed at the end of each experiment by destroying the patch and rapidly moving an open pipette tip across the liquid junction interface to measure the resulting current (10-90% rise times typically ranged between 200-500 µs).

Macroscopic recordings assessing agonists or antagonist dose-response relationships were performed using three of the four perfusion tools barrels. For half-maximal effective concentration (*EC_50_*) experiments, the centre and left barrel would contain a control and saturating glutamate solution, respectively. The right barrel would contain varying concentrations of glutamate. The receptors’ response to the different ligands was determined by alternating between the saturating pulse and the varying ligand “test” pulse and measuring the fractional evoked steady-state (I_ss_) response of the test pulse compared to the maximal I_ss_ response in the saturating pulse. Using this method, the effect of run-down (time-dependent changes in ion channel activity) was minimised.

For Schild analysis, glutamate dose response relationships were generated in the absence or presence of the competitive antagonist NBǪX. AMPA receptor desensitisation blocker cyclothiazide (CTZ, 100 μM) was included in all external solutions. Where the glutamate concentration of the test pulse exceeded 30 mM, the concentration of the control pulse was increased equivalently.

Perampanel inhibition curves were generated using all four barrels, and antagonist *IC_50_* was determined by measuring the initial peak glutamate response and comparing it to the peak evoked by an equivalent concentration of glutamate in the presence of a varying antagonist concentration.

### Data analysis

Recordings were analysed using Igor Pro 9.01 (Wavemetrics). Activation and decay times were measured using 10-90% of the slope. In cases where AMPA receptor deactivation and desensitisation displayed two exponential components, weighted time constant (τ_w, des_ and τ_w, deact_) was calculated according to:

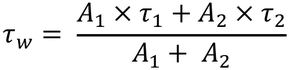

where *A_1_* and *τ_1_*, and *A_2_* and *τ_2_*, are the amplitude and time constant of the exponential components.

Statistical tests were used as described in the text. Randomisation test (non-parametric, non-paired, 10 000 iterations) was used alongside t-test as indicated as it is particularly suited for small sample numbers.

Recovery from desensitisation (*τ_rec_*) was measured using a paired-pulse protocol consisting of a desensitising glutamate application (10 mM, 200 ms, −60 mV), followed by a 10 ms test pulse delivered at interpulse intervals ranging from 3 to 300 ms. Recovery at each interval was quantified from the fractional amplitude of the test response relative to the initial peak response. The resulting recovery time course was fitted with a Hodgkin-Huxley-type function (Carbone C Plested, 2012):

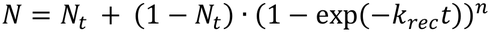

where *N* is the fraction of receptors recovered at time *t*, *N_t_* represents the fraction available immediately following the conditioning pulse, *k*_rec_ is the recovery rate constant, and *n* is the slope. The recovery time constant (τ_rec_) was derived from 1/*k_rec_*. Maximal recovery was constrained to unity during fitting. Recovery curves were compared by global nonlinear regression using an extra sum-of-squares F test.

Macroscopic current-voltage (I-V) relationships were constructed from peak current amplitudes measured over a range of holding potentials (-80 to +100 mV, in 20 mV increments). Currents were normalised to the peak response at –80 mV.

Half-maximal effective concentration (*EC*_50_) or half-maximal inhibitory concentration (*IC*_50_) values were derived by fitting the Hill equation:

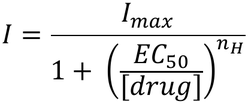

*I*_max_ denotes the peak of the fit, *EC*_50_ (or *IC*_50_ for an inhibitor) is the ligand concentration at the half-maximal response, *n*_H_ is the Hill coefficient, and [drug] refers to the equilibrium ligand concentration.

Concentration-response data were fitted with a four-parameter logistic equation with a variable slope:

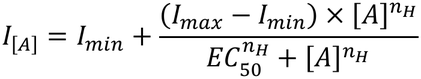

where *I*_[*A*]_ is the response at agonist concentration, [*A*]; *I*_max_ and *I*_min_ are the upper and lower asymptotes, respectively; *EC*_50_ is agonist concentration producing half-maximal response; and *n_H_* is the Hill slope.

Concentration-inhibition data were fitted with a four-parameter logistic equation with a variable slope:

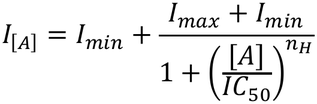

where *I*_[*A*]_ is the response at antagonist concentration [*A*]; *I*_max_ and *I*_min_ are the upper and lower asymptotes, respectively; *IC*_50_ is the antagonist concentration producing half-maximal inhibition; and *n_H_* is the Hill slope.

Schild analysis was conducted by first determining the glutamate *EC*_50_ in the absence of an antagonist. Subsequent *EC*_50_ values were measured with increasing concentrations of NBǪX (3, 10 and 30 μM). The higher the concentration of the competitive antagonist, the more agonist is needed to reach the same level of activation (as agonist and competitive antagonist are competing for the same binding site and assuming the same fraction of receptors is activated in order to generate the same response in the absence and presence of the antagonist). For each antagonist concentration, the ratio of *EC_50 with NBǪ_*_X_ over *EC_50 w/o NBǪX_* was calculated, and Schild plot was formed using the following equation:

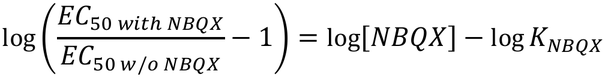

where *EC*_50_ *_with NBǪX_* is *EC*_50_ for glutamate in the presence of antagonist NBǪX at equilibrium concentration [NBǪX], *EC*_50_ *_w/o NBǪX_* is *EC*_50_ for glutamate in the absence of NBǪX and *K*_NBǪX_ the binding constant for NBǪX (for derivation of the Schild equation, see (SCHILD, 1949; Wyllie C Chen, 2007)). (*EC*_50_ *_with NBǪX_ / EC*_50_ *_w/o NBǪX_* – 1) *vs.* [NBǪX] on a logarithmic scale will appear as a straight line with a slope of 1 (Fig. 2F). At the point where *EC*_50_ *_with NBǪX_ / EC*_50_ *_w/o NBǪX_* equals 2, the corresponding concentration of the antagonist [NBǪX] on the x-axis is equal to its binding constant, *K*_NBǪX_. In this way, Schild analysis allows direct determination of *K*_NBǪX_ independently of the nature of the agonist.

Open probability (*P_o_*) from single-channel recordings of *lc* mutants was estimated using the charge passed during the channel activity as described in (Plested C Baranovic, 2016). Briefly, the charge passed by the channel during the steady-state application of glutamate was determined by integrating the area under the single-channel trace and divided by the theoretical maximum charge that would have been passed if the channel was fully open 100% of the time.

## Acknowledgements

AJE performed all patch clamp recordings and analysed the data with the help of JB and DJAW. CA performed all the cloning relevant for the project. JB designed and supervised the project. JB wrote the manuscript with the help of AJE. All authors read and contributed to the final version of the manuscript.

This work was supported by the UKRI Biotechnology and Biological Sciences Research Council (BBSRC) grant number BB/T00875X/1 (East of Scotland Biosciences Doctoral Training Partnership) to AJE, by the AMS Springboard Award (supported by Wellcome Trust, British Heart Foundation, BEIS and Diabetes UK) to JB (SBF007\100104) and CA held a Daphne Jackson Fellowship funded by Medical Research Scotland.

